# Relationship between cognitive flexibility and disordered eating attitudes across the non-clinical spectrum

**DOI:** 10.64898/2026.06.17.732879

**Authors:** Dicle Karacadag, Bianka Brezóczki, Emanuele Ciardo, Teodóra Vékony, Dezső Németh, Ana María González Martín

## Abstract

Disordered eating attitudes exist on a continuum that extends well below clinical diagnostic thresholds, yet the cognitive correlates of this non-clinical variation remain incompletely understood. Previous research linking executive functioning to disordered eating in non-clinical samples has relied almost exclusively on self-report questionnaire measures of executive function, which show weak correspondence with performance-based assessments. This methodological reliance leaves open the question of how objective executive performance relates to eating behavior across the spectrum. The present study addressed this gap by examining the association between performance-based measures of executive function and disordered eating attitudes in a non-clinical sample of 243 university students via an online experiment, using a dimensional approach consistent with the Research Domain Criteria framework. Participants completed established neurocognitive tasks covering three executive function domains: working memory was assessed with Digit Span and an N-back task, inhibitory control with a Go/No-Go task, and cognitive flexibility with the Card Sorting Task. Disordered eating attitudes were indexed using the EAT-26 total score and its subscales. A notable correlation was identified between cognitive flexibility and disordered eating attitudes, while working memory and inhibitory control exhibited no such association. Overall, our findings provide evidence for associations between executive functioning and disordered eating attitudes in a non-clinical sample.

## 1. Introduction

Given that food is a biological necessity, eating plays a fundamental role in human life. Eating behavior, however, has a complex nature, as it is shaped by emotional, motivational, cognitive, and environmental factors, and is driven by both bottom-up processes, such as the smell of food and appetite, and top-down mechanisms that guide decision-making and food choices (Beaumont, 2024). Eating behavior refers to all behavioral responses to food, including features related to the frequency and quantity of consumption, and can be conceptualized along a continuum ranging from healthy to disordered eating (Beaulieu & Blundell, 2021; Smolak & Levine, 2015). Healthy eating is characterized by a balance between the internal regulation of dietary intake in line with caloric needs and the ability to resist eating when satiated, whereas disordered eating represents the opposite end of the spectrum and includes behaviors such as restriction, binge eating, and non-nutritional consumption. Since this regulation requires both top-down and bottom-up processes (Myers, 2015), executive functions, the higher-order cognitive control processes encompassing inhibitory control, working memory, and cognitive flexibility (Diamond, 2013), are thought to play a central role. However, their influence on eating behaviors along the continuum from healthy to disordered eating is not fully understood. Uncovering this relationship in non-clinical populations is particularly important, as non-clinical variations in eating-related tendencies can still meaningfully reflect vulnerability factors without meeting diagnostic thresholds (Peschel et al., 2024). The present study, therefore, aims to examine the relationship between executive functions and maladaptive eating attitudes by assessing all three core executive domains and their associations with variation in disordered eating attitudes in a non-diagnosed university student sample.

While certain eating disorder profiles (e.g., anorexia nervosa, bulimia nervosa, and binge eating disorder) have been associated with distinct profiles of executive functioning (Eichen et al., 2017), a relatively smaller but growing body of research has extended this work to examine these associations dimensionally in non-clinical samples. This dimensional approach aligns with the Research Domain Criteria (RDoC) framework proposed by the National Institute of Mental Health (Insel et al., 2010; Cuthbert & Insel, 2013) and more recent dimensional models of psychopathology emphasizing continuous variation across diagnostic boundaries (Conway & Krueger, 2026). RDoC advocates moving beyond categorical diagnostic boundaries and instead investigating psychopathology as continuous variation in underlying neurobehavioral systems, including cognitive systems such as cognitive control and working memory, across the full range from typical to disordered functioning. Several studies have adopted this perspective, using instruments such as the Eating Attitudes Test (EAT-26; EAT-40) or the Eating Disorder Inventory (EDI-2; EDI-3) to capture variation along the eating-behavior continuum. They have reported consistent associations between subclinical disordered eating attitudes and executive functioning. Ciszewski et al. (2020) examined this relationship in a non-clinical sample of 188 university students using the EAT-26, EDI-3, and the Behavior Rating Inventory of Executive Function-Adult Version (BRIEF-A), and found that disordered eating behaviors were positively associated with self-reported problems in emotional control, shifting, inhibition, and self-monitoring. In hierarchical regression analyses, BRIEF-A emotional control emerged as the most consistent predictor of EAT-26 and EDI-3 scores, and EDI-3 bulimia scores were additionally predicted by problems with inhibition. Limbers and Young (2015) examined the relationship between BRIEF-A executive function ratings and dietary intake in university students, finding that better initiation skills were associated with higher fruit and vegetable consumption, while inhibition difficulties were associated with greater intake of saturated fat. Taken together, these results indicate that executive dysfunction is not confined to clinical eating disorders but can be detected across the non-clinical spectrum.

A critical methodological limitation is shared across this literature. Each of the studies cited above relied exclusively on self-report questionnaire measures of executive functioning, most commonly the BRIEF-A, rather than performance-based neurocognitive tasks. This stands in tension with a core methodological principle of the RDoC framework itself, which emphasizes objective, performance-based assessments of cognitive constructs over subjective self-report (Insel et al., 2010). The distinction matters, since self-report instruments such as the BRIEF-A index everyday executive behavior shaped by metacognitive insight and self-perception, whereas performance-based tasks capture momentary cognitive processing under controlled conditions, and the two methods show weak or null associations with one another (DiCarlo et al., 2026; Toplak et al., 2013). Conclusions drawn from questionnaire-based studies about the executive correlates of non-clinical disordered eating may therefore not directly reflect the underlying neurocognitive mechanisms. Based on our best knowledge, no published study has examined the relationship between EAT-26-indexed disordered eating attitudes and performance-based measures of all three core executive function domains, namely working memory, inhibitory control, and cognitive flexibility, in a non-clinical sample. Therefore, the present study was designed to address this gap.

The aim of the present study was to examine how performance-based measures of executive functioning relate to variation in disordered eating attitudes across the non-clinical spectrum. To address the methodological limitations of previous studies, we assessed all three core executive domains identified by Diamond (2013), allowing us to examine whether associations with disordered eating attitudes reflect executive functioning broadly or are specific to particular executive components in a non-clinical sample. Working memory was measured with the Digit Span (Jacobs, 1887) and N-back tasks (Owen et al., 2005). Inhibitory control was assessed using the Go/No-Go task (Miller & Low, 2001), and cognitive flexibility was measured with the Card Sorting Task (Nyhus & Barceló, 2009). Maladaptive eating attitudes were indexed with the EAT-26 (Garner & Garfinkel, 1979; Garner et al., 1982), which provides a total score along with subscales for dieting, bulimia, and food preoccupation, and oral control. The central question driving this work was whether subclinical variation in eating attitudes is associated with objectively measurable differences in executive performance and if so, which executive components carry that association.

## 2. Methods

### 2.1 Participants

A total of 344 university students aged 18-54 participated in an online study in exchange for course credits. All participants provided written informed consent and received no financial compensation. The study was approved by the Research Ethics Committee of Eötvös Loránd University (Approval No. 2021/504) and conducted in accordance with the principles of the Declaration of Helsinki.

Participants were excluded based on predefined inclusion and data quality criteria (Brezóczki et al., 2025; Hann et al., 2026; Vékony et al., 2025). Exclusion criteria included invalid age reports, self-reported clinical diagnoses of neurological disorders (e.g., epilepsy, history of head injury with loss of consciousness) and psychiatric disorders (e.g., autism spectrum disorder [ASD], attention-deficit/hyperactivity disorder [ADHD], obsessive–compulsive disorder [OCD], schizophrenia, depression, anxiety disorders, eating disorders [ED], personality disorders [PD], and post-traumatic stress disorder [PTSD]), current medication use affecting cognitive performance, recreational drug use, alcohol consumption within 6 hours before experiment, failure to complete the experimental session, and invalid task engagement failure to pass attention checks embedded in the questionnaire battery (Sarna et al., 2026; Rodd, 2024). Exclusion criteria were applied sequentially. The participant flow of the present study is shown in Figure 1. Following these exclusions, the final sample consisted of 243 participants (*M*_age_ = 22.34 years; *SD* = 4.54; 180 females, 60 males, 2 other, 1 preferred not to disclose).

**Figure 1.**
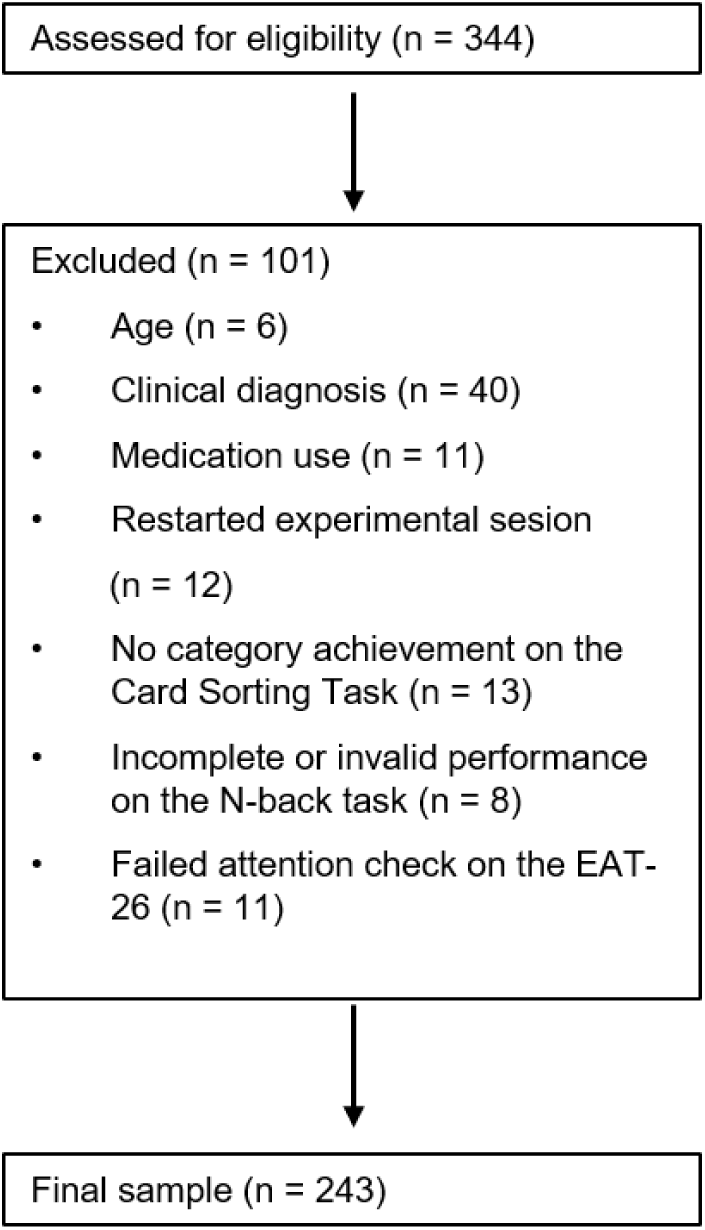
The flow diagram summarizing participant exclusions. Sequential application of exclusion criteria was used, in which each participant was assigned to a single exclusion category based on the first criterion met.

### 2.2 Measures

#### 2.2.1 Cognitive Tasks

##### 2.2.1.1 Inhibitory Control

To assess inhibitory control, a modified Go/No-Go task was administered (Bezdjian et al., 2009; Vékony, 2022). Stimuli were presented within a 2 × 2 grid, with one blue star displayed in each quadrant. On each trial, one of the stars was replaced by either the letter “P” or “R” in a random position. Participants were instructed to press the spacebar when the letter “P” appeared (Go trials) and to withhold their response when the letter “R” was presented (No-Go trials). Following the completion of half of the task, the stimulus–response mapping was reversed, such that participants were required to respond to “R” and inhibit responses to “P”. The task began with a practice phase consisting of 20 trials, during which performance feedback was provided. Subsequently, participants completed two experimental blocks of 120 trials each, with no feedback given. The ratio of Go to No-Go trials was maintained at 80:20. Each stimulus remained visible until a response was made, with a maximum response window of 500 ms. Consecutive trials were separated by a 1500 ms response-to-stimulus interval. Task performance was assessed using correct responses on Go trials, false alarms on No-Go trials, and a discriminability score (d′), with higher scores indicating greater response discriminability. Hit and false-alarm rates of 0 or 1 were adjusted using the log-linear correction before computing d′. The discriminability score was computed individually as follows:

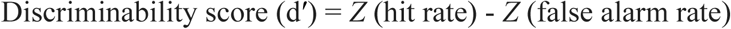

##### 2.2.1.2 Cognitive Flexibility

Cognitive flexibility was assessed using a computerized Card Sorting Task (Fox et al., 2013; Vékony, 2022) based on the Wisconsin Card Sorting Task (WCST; Berg, 1948). In this task, four cards were shown at the top of the screen, each differing in color (red, yellow, green, or blue), number of items (one to four), and shape (triangle, star, diamond, or circle). A target card appeared below these cards. Participants were asked to match the target card to one of the four cards based on color, number, or shape by clicking on their choice. The correct rule was not told to them, so they had to figure it out using the feedback given after each response. The task included 64 trials in total. The sorting rule changed after ten correct responses in a row, following a fixed order (color, shape, number, repeated twice). Performance was measured using the number of perseverative errors and the number of categories achieved. Perseverative errors refer to continuing to use the same rule even after receiving negative feedback. Categories achieved refers to the number of sorting rules correctly identified and completed. Fewer perseverative errors and more categories achieved indicate better cognitive flexibility.

##### 2.2.1.3 Working Memory

Working memory was assessed using the Digit Span task (Jacobs, 1887; Racsmány et al., 2006), reflecting verbal working memory capacity and phonological loop functioning, and a 1-back version of the N-back task (Kirchner, 1958), reflecting updating processes and continuous monitoring of information. In the 1-back task, a sequence of letters (B, K, Q, T, H, M, N, P, X, or R) was presented on the screen. Participants were required to indicate whether the currently presented letter matched the one shown immediately before it. Responses were made by pressing the “J” key for targets (same letter) and the “F” key for non-targets (different letter). The proportion of target trials was set at 20%. The task began with a 10-trial practice block with feedback, followed by 100 trials without feedback. Each stimulus was presented for 500 ms or until a response was made, with an interstimulus interval of 1500 ms. Performance was evaluated using accuracy, false alarms, reaction times, and a discriminability score calculated similarly to the Go/No-Go task.

In the Digit Span task, participants were presented with sequences of digits and were asked to recall them in the same order. The sequence length increased progressively, starting from three digits. Each sequence length included multiple trials, and participants advanced to the next level if they correctly recalled at least three sequences out of four trials. The Digit Span score was defined as the maximum sequence length correctly recalled (Conway et al., 2005).

#### 2.2.2 Self-Report Measure: The Eating Attitudes Test (EAT-26)

To assess eating-related concerns and attitudes, we used EAT-26 self-report questionnaire (Garner & Garfinkel, 1979; Garner et al., 1982), a widely used screening tool derived from the EAT-40, validated in Hungarian samples by Túry et al. (1990). The scale consists of 26 items rated on a 6-point Likert scale (Always to Never), with higher scores indicating greater levels of eating-related concerns. Example items include “I am terrified about being overweight” and “I find myself preoccupied with food.” The EAT-26 provides a total score reflecting overall disordered eating attitudes, as well as three subscale scores: Dieting, Bulimia and Food Preoccupation, and Oral Control. A total score of 20 or above is generally considered indicative of elevated risk and may warrant further clinical evaluation (Garner et al., 1982). The Dieting subscale captures restrictive eating and weight concern, the Bulimia and Food Preoccupation subscale reflects thoughts about food and binge-related tendencies, and the Oral Control subscale assesses self-control around eating and perceived social pressure related to food intake.

### 2.3 Procedure

Data were collected online using the Gorilla Experiment Builder platform (https://www.gorilla.sc/) (Anwyl-Irvine et al., 2020). Participants completed the study in a single session consisting of computerized cognitive tasks and self-report questionnaires, using either a desktop computer or a laptop. To minimize potential order effects, the four cognitive tasks were administered in a randomized sequence across participants. In addition, demographic information, including socioeconomic status, sleep characteristics, health status, and handedness, was collected. To account for potential confounding factors associated with the online testing environment, participants provided brief information about their testing conditions; these details are described in the ‘Participants’ section. The experimental tasks were implemented using the jsPsych library (de Leeuw, 2015). The source code for all tasks is openly available via the link provided in the Data Availability Statement.

### 2.4 Statistical Analysis

All data processing, statistical analyses, and visualization were conducted using R version 4.5.1 (R Core Team, 2025). Data manipulation and visualization were performed using packages from the *tidyverse* (Wickham et al., 2019), and descriptive statistics were computed using the *psych* package (Revelle, 2024).

Prior to analysis, the dataset was screened for missing values, distributional assumptions, and univariate outliers (z-scores > |3|). Gender was coded as a categorical variable (reference-coded factor) with female as the reference category. Outliers were retained to preserve the natural variability of the sample and to avoid excluding potentially meaningful individual differences. Normality was assessed using the Shapiro–Wilk test in conjunction with skewness and kurtosis indices. Following the application of exclusion criteria, no further missing values were present in the final dataset, and all analyses were conducted on the same sample (N = 243).

Most study variables deviated from normality. Specifically, all outcome variables (EAT-26 total score and subscale scores; see Figure 2) and most executive function predictors (Card Sorting Task category achievement, perseverative errors, N-back d′, and Digit Span scores) did not meet normality assumptions, whereas Go/No-Go d′ scores did not significantly differ from a normal distribution (p = .09). Based on these distributional characteristics, we fitted a Generalized Linear Model (GLM) to describe the relationship between the outcome variables and the predictors. GLMs were estimated using functions implemented in the base R stats package. Regression coefficients, standard errors, test statistics, and associated p-values are reported. Statistical significance was evaluated using two-tailed tests with α set at .05.

**Figure 2.**
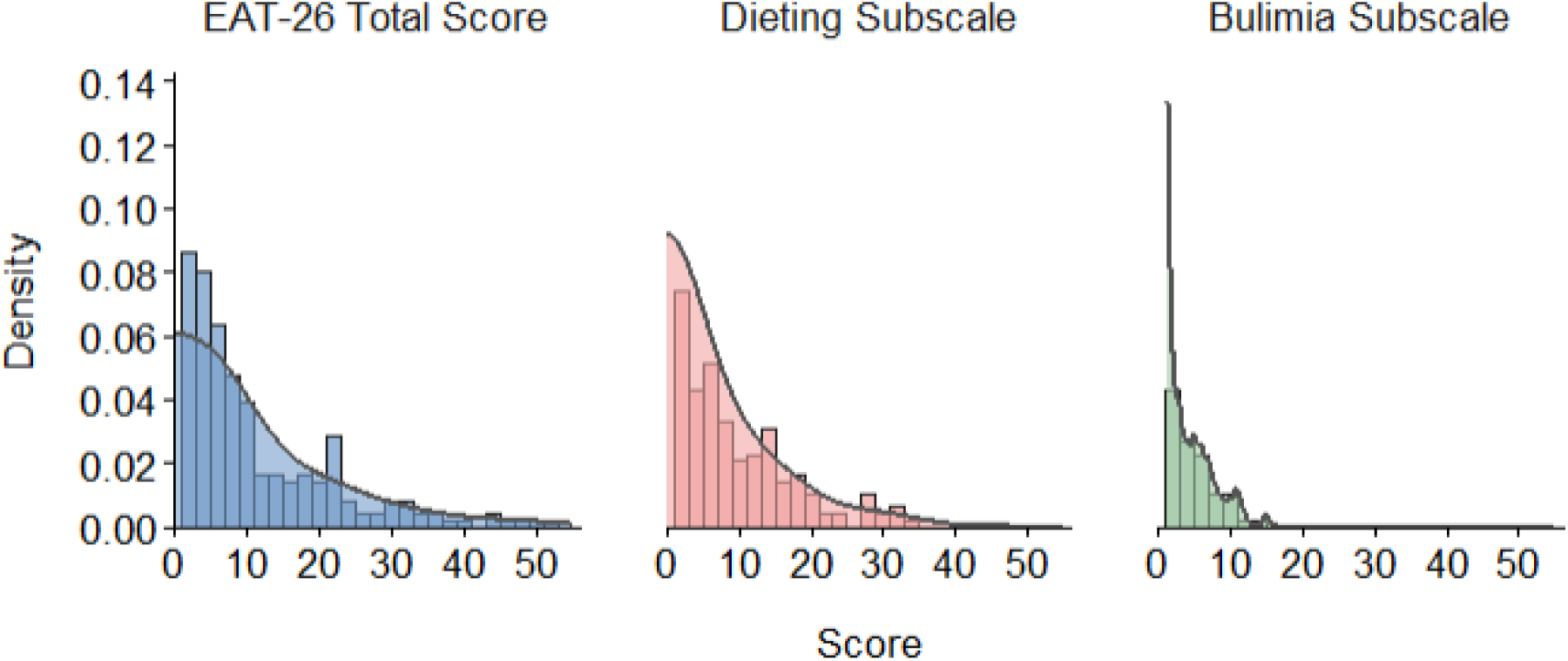
Density distributions and zero accumulation profiles for EAT-26 total and subscale scores. The multi-panel visualization displays the relative frequency of scores via histogram bins, overlaid with smoothed kernel density curves. First two panels (EAT-26 Total Score and Dieting) exhibit a prominent right-skewness (positive skew) and last graph shows the bulimia subscale exhibiting a pronounced floor effect, with a substantial proportion of scores at zero. These distributional features support the use of Gamma regression models with a log link on (outcome + 1) transformed variables.

Measurements derived from neurobehavioral tasks, including Digit Span, Go/No-Go d′, N-back d′, and Card Sorting Task indices, were included as predictors. Separate models were estimated for each eating attitude outcome (EAT-26 total score and subscale scores), while controlling for gender (female, male, other) and age. To accommodate positive skewness, models predicting the EAT-26 total score, dieting, and bulimia subscales were initially specified using a Gamma distribution with a log link function. A constant of 1 was added to all outcome variables prior to model estimation to ensure strictly positive values, as required for Gamma regression given the presence of zero scores in the subscales (i.e., outcome + 1 transformation).To identify the most parsimonious model and select the most relevant predictors, a bidirectional stepwise model selection procedure (both forward selection and backward elimination) was applied to the initial GLMs. This selection was conducted using the step() function in base R, which evaluates model combinations by minimizing the Akaike Information Criterion (AIC). Only the predictors retained in the final, optimized models were interpreted.

## 3. Results

### 3.1 Descriptive Statistics of Study Variables

Descriptive statistics for all study variables are presented in Table 1. Variables showed deviations from normality, as indicated by skewness and kurtosis values, which informed subsequent model selection.

**Table 1.** Descriptive Statistics of Study Variables.

| Variable | N | M | SD | Median | Min | Max | Skewness | Kurtosis |
| --- | --- | --- | --- | --- | --- | --- | --- | --- |
| Age (years) | 243 | 22.34 | 4.54 | 21.00 | 18.00 | 54.00 | 3.69 | 17.39 |
| EAT-26 Total Score | 243 | 11.47 | 10.52 | 8.00 | 0.00 | 52.00 | 1.57 | 2.19 |
| Dieting | 243 | 7.27 | 7.91 | 5.00 | 0.00 | 34.00 | 1.35 | 1.34 |
| Bulimia/Food Preoccupation | 243 | 1.42 | 2.65 | 0.00 | 0.00 | 15.00 | 2.39 | 5.72 |
| Oral Control | 243 | 2.78 | 2.45 | 2.00 | 0.00 | 16.00 | 1.63 | 4.36 |
| CST Perseverative Errors | 243 | 8.33 | 3.97 | 7.00 | 1.00 | 23.00 | 1.48 | 2.08 |
| CST Categories Achieved | 243 | 3.57 | 1.07 | 4.00 | 1.00 | 5.00 | -0.77 | 0.00 |
| Digit Span | 243 | 5.98 | 1.32 | 6.00 | 3.00 | 9.00 | 0.45 | -0.22 |
| N-back d' | 243 | 2.46 | 1.31 | 2.76 | -2.80 | 4.14 | -1.52 | 2.41 |
| Go/No-Go d' | 243 | 1.31 | 0.64 | 1.30 | -0.18 | 3.31 | 0.31 | -0.18 |

**Note.** N = sample size; M = mean; SD = standard deviation. Skewness and kurtosis values are reported to indicate distributional characteristics.

### 3.2 Preliminary Psychometric Analyses

Prior to the main analyses, the internal consistency of the EAT-26 total score and its subscales was examined to evaluate which EAT-26 subscales could be included. The total score and the dieting subscale demonstrated good internal consistency (Cronbach’s α = .89 for both), whereas the bulimia subscale showed acceptable reliability (Cronbach’s α = .79). In contrast, the oral control subscale demonstrated unacceptable internal consistency (α = .45), falling substantially below conventional thresholds for research use. Consequently, this subscale was excluded from subsequent analyses.

### 3.3 Relationship Between Executive Functions and Eating Attitudes

The analyses examined the associations between executive function measures and disordered eating attitudes across three outcome variables: EAT-26 total score and its dieting and bulimia subscales. For each outcome, age, gender, and executive function measures were included as predictors. Based on these findings, GLMs with a Gamma distribution and log link function were employed as the primary inferential approach for all outcome variables.

#### 3.3.1 EAT-26 Total Score

A likelihood ratio test comparing the final step-wise selected model to the intercept-only model indicated that the selected model significantly improved fit, χ²(5) = 18.27, *p* = .003, suggesting that the retained set of demographic and executive function predictors was collectively associated with EAT-26 total scores.

Examination of the parameter estimates revealed that gender and age significantly predicted EAT-26 total scores. Relative to female participants (reference category), male participants showed significantly lower expected EAT-26 scores (β = −0.412, SE = 0.119, t = −3.477, *p* < .001), corresponding to an approximate 34% reduction in expected scores on the response scale. Age emerged marginally as a negative predictor, indicating that older age was associated with slightly lower EAT-26 scores (β = −0.023, SE = 0.011, t = −2.014, *p* = .045).

Following the stepwise selection procedure, only two executive function measures were retained in the final model. Cognitive flexibility, as measured by CST perseverative errors, showed a robust positive association with EAT-26 scores, indicating that a higher number of perseverative errors was significantly associated with higher expected disordered eating attitudes (β = 0.037, SE = 0.013, t = 2.862, *p* = .005). Working memory performance, indexed by N-back d’, was retained but demonstrated only a marginal negative association with EAT-26 scores (β = -0.070, SE = 0.040, t = -1.755, *p* = .081). The forest plot provides a visual summary of regression coefficients and confidence intervals for executive function predictors across all eating attitude outcomes (see Figure 3).

**Figure 3.**
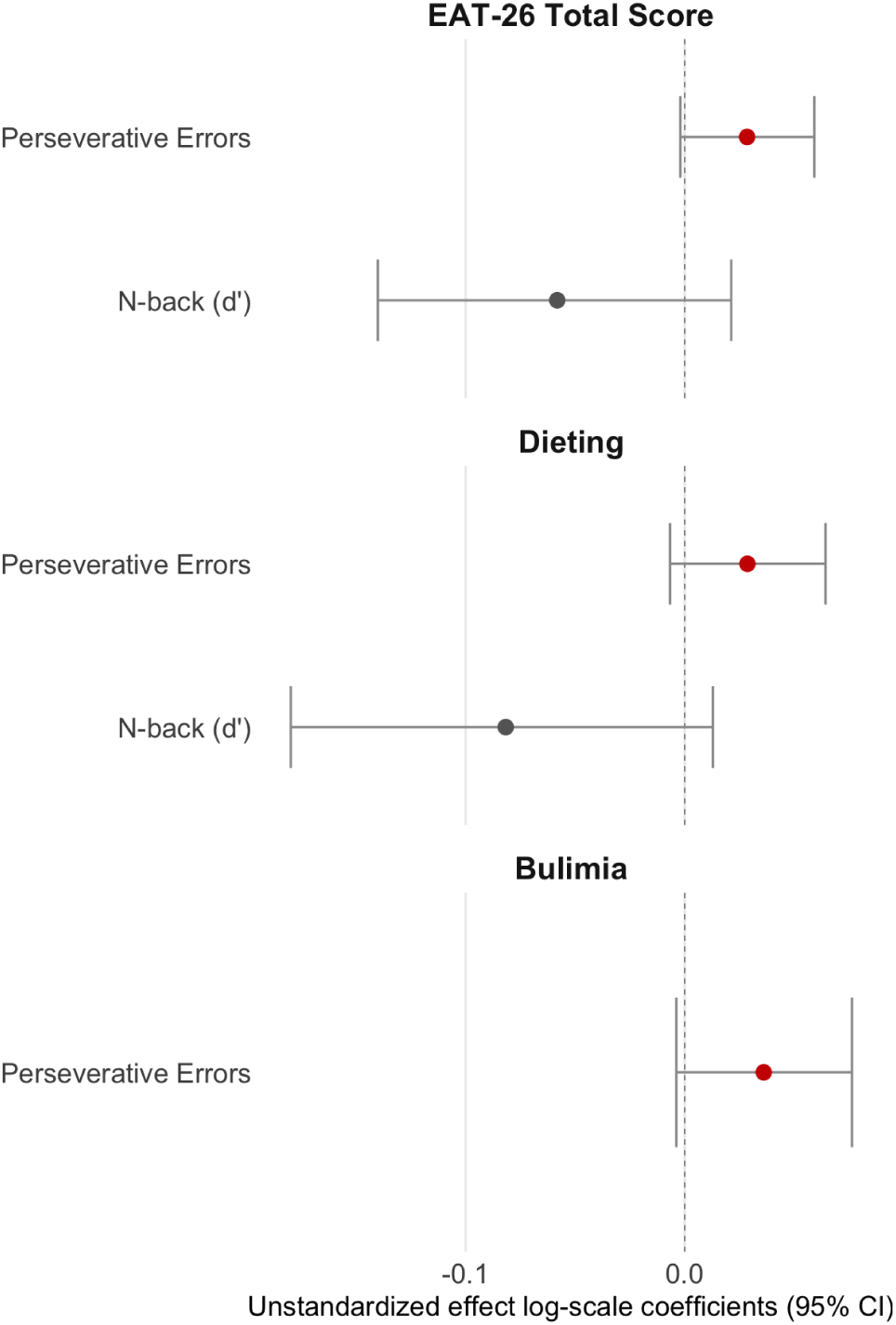
Forest plot of associations between executive function measures and disordered eating attitudes across EAT-26 total score, dieting, and bulimia subscale models. Unstandardized effect log-scale coefficients and 95% confidence intervals are displayed for the retained predictors. Positive values indicate higher disordered eating attitudes, whereas negative values indicate lower scores. It is important to note that for perseverative errors, elevated scores indicate poorer performance, whereas for the other measures, elevated scores signify better performance. The vertical line at zero represents the null effect.

#### 3.3.2 Dieting Subscale

A likelihood ratio test comparing the stepwise-selected model to the intercept-only model indicated that the overall model significantly improved fit, χ²(4) = 23.57, *p* < .001, suggesting that the retained set of demographic and executive function predictors was significantly associated with dieting-related eating attitudes.

Examination of the parameter estimates revealed that gender significantly predicted dieting scores. Relative to female participants (reference category), male participants showed significantly lower dieting scores (β = −0.565, SE = 0.140, *t* = −4.034, *p* < .001). Following the stepwise selection procedure, only two executive function measures were retained in the final model. Age, Digit Span, CST category achievement, and Go/No-Go d’ were eliminated. Cognitive flexibility, as measured by CST perseverative errors, showed a significant positive association with dieting scores (β = 0.032, SE = 0.016, *t* = 2.120, *p* = .035), indicating that higher perseverative error scores were associated with higher expected dieting attitudes. Working memory performance, indexed by N-back d’, was retained but did not reach significance (β = −0.075, SE = 0.046, t = −1.640, *p* = .102).

#### 3.3.3 Bulimia and Food Preoccupation Subscale

A likelihood ratio test comparing the stepwise-selected model to the intercept-only model indicated that the overall model significantly improved fit, χ²(4) = 22.15, *p* < .001, suggesting that the retained set of demographic and executive function predictors was significantly associated with bulimia-related eating symptoms.

Examination of the parameter estimates showed that gender significantly predicted bulimia scores. Relative to the female participants (reference category) male participants exhibited significantly lower bulimia scores (β = −0.406, SE = 0.148, *t* = −2.738, *p* = .007). Age was retained by the selection procedure but demonstrated only a marginal negative association with bulimia scores (β = -0.023, SE = 0.014, *t* = -1.662, *p* = .098). Only one executive function measure was retained in the final model: Cognitive flexibility, measured by the CST perseverative errors, showed a highly significant positive association with bulimia scores (β = 0.051, SE = 0.016, *t* = 3.186, *p* = .002), indicating that the higher number of perseverative errors was strongly associated with higher expected bulimia symptoms. Digit Span, CST category achievement, Go/No-Go d′, and N-back d′ were eliminated from the model (see Figure 3).

## 4. Discussion

Our study examined how performance-based measures of executive functioning relate to maladaptive eating attitudes in a non-clinical sample within a dimensional psychopathology framework. The findings showed that cognitive flexibility, indexed by perseverative errors on the Card Sorting Task, was associated with EAT-26 total scores and both dieting and bulimia subscales. While N-back d′ was retained in the model of the total score and the dieting subscale, it was not significantly related to disordered eating attitudes. No additional cognitive functions were preserved in the models. Taken together, these findings suggest that cognitive flexibility, but not other measures, was consistently associated with maladaptive eating attitudes in this non-clinical sample. However, the effects involving cognitive flexibility observed across multiple eating-related outcomes may indicate that its association is not specific to particular symptom subtypes, but rather reflects a broader relationship with general maladaptive eating attitudes.

Given the fact that previous non-clinical studies using the EAT-26 have relied mainly on self-report measures of executive functioning, rather than performance-based cognitive tasks, our findings are not straightforward to compare directly with earlier work. They sit between two literature bodies: clinical studies using neurocognitive tasks (Favieri et al., 2024; Weinbach et al.,2020; Allom & Mullan, 2014), and non-clinical studies using questionnaires about everyday executive difficulties (Limbers & Young, 2015; İçen et al., 2025; Ciszewski et al.,2020). In clinical samples, different eating disorder profiles have often been linked to partly different cognitive mechanisms, although the picture is far from settled. Restrictive presentations of anorexia nervosa, for example, are commonly associated with rigid behavioural patterns, persistent dieting, and compulsive weight control. These features have been linked to reduced cognitive flexibility and may also contribute to treatment resistance (Krug et al., 2024; Stedal et al., 2021). By contrast, binge-related conditions, including bulimia nervosa, binge eating disorder, and the binge–purge subtype of anorexia nervosa, have more often been associated with impulsivity and impaired inhibitory control, which fits with recurrent episodes of uncontrolled eating (Krug et al., 2024).

Still, findings from clinical groups cannot be mapped directly onto non-clinical eating attitudes. Most clinical studies compare diagnosed patients with undiagnosed controls. This captures the two ends of the continuum but says less about what happens between them. Evidence from non-clinical samples is also limited and inconsistent. For example, İçen et al. (2025) reported an association only between dieting scores and executive functioning, whereas Ciszewski et al. (2020) found positive associations between EAT-26 total, dieting, and bulimia scores and self-reported executive difficulties, including inhibition and set-shifting. The present study adds to this literature by examining these associations using performance-based measures across all three core executive function domains. A significant association was observed between cognitive flexibility and overall disordered eating attitudes indexed by the EAT-26 total score. The presence of significant findings involving cognitive flexibility across the subscales may suggest that this relationship is not confined to a particular eating-related symptom profile, although this possibility requires further investigation.

Eating behavior is inherently dynamic and requires the continuous integration of competing signals, including physiological states, such as hunger and satiety, external food cues, and internally generated cognitive rules, such as dietary restrictions or “safe food” categorizations (Meule & Vögele, 2013). Within this framework, maladaptive eating tendencies may be driven less by global executive dysfunction than by difficulty updating or overriding rigid, rule-based behavioural strategies. Cognitive flexibility is closely related to such processes, as it supports the ability to shift away from established rules, adapt to changing internal and external demands, and modify behaviour when previous strategies are no longer appropriate. Difficulties in this domain may therefore contribute to patterns such as compulsive weight control, ritualized food consumption, and resistance to new eating habits. Although the present study was not designed to examine neural mechanisms directly, the findings can be interpreted in light of neurocognitive models of cognitive flexibility. In the present study, the relationship between cognitive flexibility and disordered eating attitudes was examined using generalized linear models. Results suggest that cognitive flexibility may represent a cognitive domain of interest in relation to overall disordered eating attitudes.

Cognitive flexibility has been closely linked to prefrontal–striatal circuits involved in behavioural adaptation, reward learning, and habit regulation (Diamond, 2013; Miller & Cohen, 2001). From this perspective, our results are consistent with the idea that prefrontal networks may play a role in the regulation of eating behaviour, particularly when this regulation requires updating food-related rules, overriding habitual responses, or adapting to changing internal and external cues. Reduced flexibility may therefore make it more difficult to override habitual eating patterns, shift away from rigid food-related rules, or disengage from persistent food-related thoughts. These processes are highly relevant to disordered eating attitudes, particularly when eating behaviour becomes organized around inflexible rules, repetitive concerns, or difficulty adapting to changing internal and external cues. Clinical neuroimaging findings are broadly consistent with this interpretation, showing alterations in prefrontal regions implicated in cognitive control and flexibility across the eating disorder spectrum (Uher et al., 2004). This neural interpretation should therefore be treated as a framework for future research rather than as direct evidence from the present study. These prefrontal systems are also critically involved in the development and maintenance of rigid behavioural patterns, which represent a core feature of disordered eating across diagnostic categories. Importantly, such rigidity-related mechanisms may already be detectable at subclinical levels, where individuals vary in their cognitive control strategies despite not meeting diagnostic thresholds. This may help explain why cognitive flexibility emerged as a consistent predictor across eating behaviour outcomes in the present sample, suggesting that it reflects a transdiagnostic and dimensional vulnerability rather than a process specific to particular symptom profiles.

The observed effect of cognitive flexibility, together with the absence of significant associations between working memory and inhibitory control and disordered eating attitudes, may indicate that these executive domains are more strongly engaged in clinically more severe presentations or under conditions of increased emotional salience, rather than in subclinical variation. The absence of significant associations may also be partly attributable to task-specific characteristics. For instance, the Digit Span task employed in the present study involved forward recall only, which is typically considered a measure of short-term verbal storage rather than a more demanding form of executive working memory. Similarly, the 1-back task may place relatively low demands on executive control and may therefore be susceptible to ceiling effects in university samples. If performance was clustered at the upper end of the distribution, this restricted variability could have further reduced the likelihood of detecting associations with eating-related outcomes.

In addition, interpretation of the findings is constrained by the task impurity problem commonly observed in executive function research (Smith et al., 2018). Although the neurobehavioural tasks employed in the present study were selected to represent the core executive function domains (Diamond, 2013), performance on these tasks is not exclusively determined by the targeted executive process. For example, successful performance on the Card Sorting Task, commonly used as a measure of cognitive flexibility, also relies on working memory, error monitoring, feedback processing, and strategy maintenance (Gamboz et al., 2009). Similarly, performance on Go/No-Go tasks may reflect sustained attention and response speed in addition to inhibitory control (Swick et al., 2011). Therefore, the observed associations cannot be attributed exclusively to a single executive function domain. Future studies may benefit from employing multiple neurobehavioural measures for each executive function construct or using latent-variable approaches to better isolate the cognitive processes of interest. Furthermore, the tasks employed in the present study primarily assessed “cold” executive functions within an online experimental setting, which may not adequately capture the affectively and motivationally driven processes that characterize eating-related decision-making. Additionally, relatively restricted variability in executive performance within a non-clinical sample may have further limited the detection of subtle associations.

Despite these methodological considerations, the overall pattern of findings remains informative. Among the executive function measures examined, cognitive flexibility was the only domain that showed consistent associations with eating-related outcomes, whereas working memory and inhibitory control were unrelated to eating attitudes and behaviours. Although these findings should be interpreted in light of the limitations discussed above, they nevertheless provide preliminary support for the possibility that cognitive flexibility may play a particularly relevant role in non-clinical disordered eating. From a broader theoretical perspective, these findings align with dimensional frameworks and spectrum approaches such as the RDoC, which emphasize the identification of specific cognitive mechanisms rather than assuming uniform contributions of broad executive constructs (Insel et al., 2010). The present results suggest that cognitive flexibility, rather than executive functioning as a unitary system, may represent a more proximal cognitive correlate of subclinical disordered eating. This perspective may also help reconcile inconsistencies in the literature, particularly in studies relying on self-report measures of executive functioning, which tend to capture perceived everyday difficulties rather than performance-based cognitive processes.

A few methodological considerations should be taken into account when interpreting these findings and may guide future research. Since the present study used a cross-sectional design, it cannot determine causality or the direction of the association between executive functioning and eating-related attitudes. Future longitudinal and experimental studies will therefore be important to clarify whether executive difficulties contribute to maladaptive eating patterns, whether sustained eating patterns gradually shape executive functioning, or whether both processes interact over time. In addition, our non-clinical sample provides a useful dimensional perspective, but future work should examine whether similar associations are observed in clinical populations, where executive difficulties and eating-related symptoms may be more pronounced. The limitations of self-report assessment should also be acknowledged. Participants were not clinically interviewed, and therefore, the presence of undiagnosed eating pathology cannot be ruled out. As a result, some individuals classified within the non-clinical range may still have exhibited clinically relevant symptoms that were not captured by the self-report measures. Although performance-based tasks offer objective indices of cognitive functioning, they may not fully capture the complexity of real-world executive demands, especially in emotionally salient or food-related contexts. Relatedly, future studies could benefit from incorporating potential confounding and moderating factors, including depression, anxiety, comorbid symptoms, and biological or developmental variables. Finally, the present findings should be interpreted in light of task-related variability. Different neurocognitive tasks may emphasize partly different processes, and the use of emotionally salient or food-related stimuli may influence the magnitude of observed effects. For example, inhibitory control difficulties are often more apparent in Go/No-Go paradigms than in tasks such as the Stroop, while findings on set-shifting may vary depending on whether measures such as the Wisconsin Card Sorting Task or the Trail Making Test are used (Smith et al., 2018). Future studies using longitudinal designs and multiple task designs would help determine how stable these executive profiles are over time and how strongly they depend on the specific cognitive demands of the task.

In conclusion, the present findings suggest that performance-based cognitive flexibility is selectively associated with maladaptive eating attitudes across the non-clinical spectrum, whereas working memory and inhibitory control do not show significant associations. By combining objective neurocognitive tasks assessing the three core executive function domains with generalized linear modeling in a non-clinical sample, the present study extends previous research that has largely relied on self-report measures of executive functioning. Furthermore, the observed linear relationship between cognitive flexibility and eating attitudes was examined using generalized linear models, supporting a stable relationship between these constructs. Together, these results converge with RDoC-aligned, transdiagnostic accounts in which difficulty updating rigid, rule-based strategies, rather than global executive dysfunction, constitutes a candidate cognitive vulnerability shared across the eating-behaviour continuum, detectable well below diagnostic thresholds. Beyond its theoretical contribution, the study has practical implications for early identification and prevention in non-clinical populations, suggesting that flexibility-targeted cognitive and behavioural interventions may offer a useful point of intervention before symptoms consolidate. Future work integrating longitudinal, multimodal (behavioural, neural, and gut–brain) designs will be needed to establish directionality and to test whether enhancing cognitive flexibility can attenuate the developmental trajectory from subclinical eating concerns to clinically significant disorder (Smith et al., 2018; Lane et al., 2023).

## Data availability

The behavioral datasets generated and analyzed during the current study, as well as the custom analysis code, are available in the OSF repository at https://osf.io/eurt9/overview?view_only=59adff1395e94560a9792b0d5a38c6da. The experiment was not preregistered.

## Declaration on the Use of AI-Assisted Technologies

The authors, who are not native English speakers, used an AI-assisted tool to improve the readability of author-written text. The tool was not used to generate scientific content or ideas. All concepts, analyses, and conclusions are the authors’ own, and the authors take full responsibility for the final manuscript.

## Contributions

D.K.: Conceptualization, Formal analysis, Writing – Original Draft, Writing – Review & Editing, Visualization

B.B: Conceptualization, Software, Validation, Investigation, Data Curation, Writing – Review & Editing

T.V: Conceptualization, Software, Validation, Investigation, Writing – Review & Editing, Supervision

E.C.: Writing – Review & Editing

D.N: Conceptualization, Software, Validation, Investigation, Writing – Review & Editing, Supervision, Funding

A.M.G.M: Conceptualization, Writing – Review & Editing, Supervision

